# Correlation and co-localization of QTL for stomatal density and canopy temperature under drought stress in Setaria

**DOI:** 10.1101/2020.10.14.339580

**Authors:** Parthiban Thathapalli Prakash, Darshi Banan, Rachel E. Paul, Maximilian J. Feldman, Dan Xie, Luke Freyfogle, Ivan Baxter, Andrew D.B. Leakey

## Abstract

Mechanistic modeling indicates that stomatal conductance could be reduced to improve water use efficiency (WUE) in C_4_ crops. Genetic variation in stomatal density and canopy temperature was evaluated in the model C_4_ genus, Setaria. Recombinant inbred lines (RIL) derived from a *Setaria italica* x *Setaria viridis* cross were grown with ample or limiting water supply under field conditions in Illinois. An optical profilometer was used to rapidly assess stomatal patterning and canopy temperature was measured using infrared imaging. Stomatal density and canopy temperature were positively correlated but both were negatively correlated with total above-ground biomass. These trait relationships suggest a likely interaction between stomatal density and the other drivers of water use such as stomatal size and aperture. Multiple QTLs were identified for stomatal density and canopy temperature, including co-located QTLs on chromosomes 5 and 9. The direction of the additive effect of these QTLs on chromosome 5 and 9 were in accordance with the positive phenotypic relationship between these two traits. This suggests a common genetic architecture between stomatal patterning in the greenhouse and canopy transpiration in the field, while highlighting the potential of setaria as a model to understand the physiology and genetics of WUE in C4 species.

**Highlight:** This article reports a phenotypic and genetic relationship between two water use related traits operating at leaf level and canopy level in a C_4_ model crop species.

## Introduction

Drought stress is the primary limiting factor to crop production worldwide (Boyer, 1982). This is underpinned by the unavoidable loss of water vapor from leaves, via stomata, to the atmosphere in order for CO_2_ to move in the reverse direction and be assimilated through photosynthesis. In the coming decades, crops are likely to experience increasingly erratic rainfall patterns, with more frequent and intense droughts, due to climate change (Stocker *et al.*, 2013). Irrigation of crops already accounts for ~70% of freshwater use, limiting the sustainability of any increase in irrigation to address drought limitations (Hamdy *et al.*, 2003). Consequently, there is great interest in understanding and improving crop water-use efficiency (WUE; Leakey *et al.*, 2019) as well as crop drought resistance (Cattivelli *et al.*, 2008).

Substantial advances have been made in understanding WUE and drought resistance at the genetic, molecular, biochemical and physiological levels in the model species, *Arabidopsis thaliana* (Zhang *et al.*, 2004; Valliyodan and Nguyen, 2006; Nakashima *et al.*, 2012). Unfortunately, efforts to translate this knowledge into improved performance of crop plants in the production environment have not resulted in success as frequently as hoped (e.g. Nelson *et al.*, 2007; Nemali *et al.*, 2015). Physiological, agronomic and breeding studies directly in crops have also resulted in improved drought avoidance and drought tolerance (e.g. Condon *et al.*, 2004; Sinclair *et al.*, 2017), but there are challenges associated with trying to apply modern systems biology and bioengineering tools to crops that are relatively large in stature and have generation times of several months. Consequently, *Setaria viridis* (L.) has been proposed as a model C_4_ grass that has characteristics that make it tractable for systems and synthetic biology while also being closely related to key C_4_ crops, so that discoveries are more likely to translate to production crops (Brutnell *et al.*, 2010; Li and Brutnell, 2011). This study aimed to assess natural genetic variation in Setaria for traits two key traits related to WUE and drought response: stomatal density and canopy temperature (as a proxy for the rate of whole-plant water use).

*Setaria italica and Setaria viridis* are model C_4_ grasses belonging to the panicoideae subfamily, which also includes maize, sorghum, sugarcane, miscanthus and switchgrass (Brutnell *et al.*, 2010; Li and Brutnell, 2011). Foxtail millet (*Setaria italica*) is also a food crop in China and India (Devos *et al.*, 1998). The availability of sequence data for its relatively small diploid (2n = 18) genome, short life cycle, small stature, high seed production, and amenability for transformation makes Setaria a good model species for genetic engineering (Brutnell *et al.*, 2010; Bennetzen *et al.*, 2012). In addition, Setaria is adapted to arid conditions and is a potential source of genes conferring WUE and drought resistance.

Whole plant WUE is the ratio of plant biomass accumulated to the amount of water used over the growing season (Condon *et al.*, 2004; Morison *et al.*, 2007; Blum, 2009; Tardieu, 2013). WUE at the leaf level is a complex trait controlled by factors including photosynthetic metabolism, stomatal characteristics, mesophyll conductance and hydraulics (Farquhar *et al.*, 1989; Condon *et al.*, 2002; Hetherington and Woodward, 2003). At the whole-plant scale it is modified by canopy architecture and root structure and function (Martre *et al.*, 2001; White and Snow, 2012).

Stomata regulate the exchange of water and carbon dioxide (CO_2_) between the internal leaf airspace and the atmosphere (Hetherington and Woodward, 2003; Bertolino *et al.*, 2019). Stomatal conductance (g_s_), which is the inverse of the resistance to CO_2_ uptake and water loss, is controlled by a combination of stomatal density, patterning across the leaf surface, maximum pore size, and operating aperture (Faralli *et al.*, 2019; Nunes *et al.*, 2020). Of these traits, stomatal density is most simple to measure (Dow and Bergmann, 2014). Consequently, genetic variation in stomatal density has been explored in a range of species, including the identification of quantitative trait loci (QTL) in rice (Laza *et al.*, 2010), wheat (Schoppach *et al.*, 2016; Shahinnia *et al.*, 2016), barley (Liu *et al.*, 2017), Arabidopsis (Dittberner *et al.*, 2018; Delgado *et al.*, 2019), brassica (Hall *et al.*, 2005), poplar (Dillen *et al.*, 2008) and oak (Gailing *et al.*, 2008). However, there is a notable knowledge gap regarding genetic variation in stomatal density within C_4_ species. While many genes involved in the regulation of stomatal development are known in Arabidopsis, investigation of whether their orthologs retain the same function in grasses and other phylogenetic groups that include the major crops is still relatively nascent (e.g. Raissig *et al.*, 2017; Lu *et al.*, 2019; Mohammed *et al.*, 2019). This is in part because standard protocols for measuring stomatal density are still laborious and time consuming, which slows the application of quantitative, forward, and reverse genetics approaches to identifying candidate genes and confirming their function. Therefore, improved methods for acquiring and analyzing images of stomatal guard cell complexes and other cell types in the epidermis are an area of active research (Haus *et al.*, 2015; Dittberner *et al.*, 2018; Fetter *et al.*, 2019; Li *et al.*, 2019). In addition, alternative approaches to rapidly screen stomatal conductance or rates of transpiration at the leaf and canopy scales (including temperature as a proxy) have also been developed and used to reveal genetic variation in traits related to drought stress and WUE (Liu *et al.*, 2011; Bennett *et al.*, 2012; Awika *et al.*, 2017; Prado *et al.*, 2018; Deery *et al.*, 2019; Vialet-Chabrand and Lawson, 2019). However, the expected links between genetic variation in stomatal density and measures of water use, which would be expected in theory, are rarely tested and when tested, the results are inconsistent (e.g. Fischer *et al.*, 1998; Ohsumi *et al.*, 2007; Kholová *et al.*, 2010; Schoppach *et al.*, 2016).

To address these questions, we used a field study of a biparental mapping population developed from an interspecific cross between *Setaria viridis* (A10) and *Setaria italica* (B100).

The study was designed with the aim of (i) applying rapid, image-based methods for phenotyping stomatal density and canopy water use; (ii) Identifying variation in stomatal patterning, canopy temperature and productivity; (iii) assessing trait relationships between stomatal density, canopy temperature and biomass production; and (iv) identifying quantitative trait loci for these traits in Setaria, grown in the field under wet and dry treatments.

## Materials and methods

### Plant material

This study used a population of 120 F_7_ recombinant inbred lines (RIL), which were generated by an interspecific cross between domesticated *Setaria italica* accession B100 and a wild-type *Setaria viridis* accession A10 (Devos *et al.*, 1998; Wang *et al.*, 1998).

### Greenhouse experiment

Variation in stomatal density among the RILs was assessed in a greenhouse study at the University of Illinois, Urbana Champaign in 2015. Plants were grown in pots (10 × 10 × 8.75 cm) filled with potting mixture (Metro-Mix 360 plus, Sun Gro Horticulture). Three seeds were sown directly into the pot. After germination, plants were thinned to one plant per pot. Growth conditions were 30/24 °C during the day/night and plants received supplemental photosynthetically active radiation from high-pressure sodium and metal halide lamps during the day (350 μmol m^−2^ s^−1^ on a 16-h day / 8-h night cycle). Throughout the growing period, water was added to pot capacity along with fertilizer (EXCEL-CAL-MAG 15-5-5) 2-3 times a week.

The youngest fully expanded leaf was excised from the plant 17 - 22 days after sowing, covered in wet paper towel, sealed in airtight bags, and stored at 4°C. Within 48 hours, a sample was excised with a razor blade from midway along the leaf to provide a cross-section from one leaf margin to the midrib (approximately 20-30 mm length, 3-20 mm wide). This sample was attached to a glass microscope slide using double-sided adhesive tape and the abaxial surface immediately imaged using an μsurf explorer optical topometer (Nanofocus, Oberhausen, Germany (Haus *et al.*, 2015). Four fields of view in a transect from the midrib to the edge of a single leaf were imaged using a 20x magnification objective lens. The images were then exported into TIF files and the stomatal number was counted using the cell counter tool in ImageJ software (http://rsbweb.nih.gov/ij/). Stomatal density was calculated by normalizing the number of stomata with the area of the field of view (0.64 mm^2^). Data from each of the four fields of view were treated as subsamples and averaged to estimate mean stomatal density for each replicate plant of a given RIL.

### Field experiment

The field experiment to assess variation in canopy temperature and total above-ground biomass was conducted at the SoyFACE field site, University of Illinois, Urbana Champaign in 2015, in the manner described by Feldman *et al.* (2017). The average air temperature over the growing season was 21.5 °C with a relative humidity of 82 % (Figure 1). In brief, plants were germinated in plug trays in the greenhouse and then after 9 days after sowing, seedlings were hand transplanted (July 15, 2015) into plots at the field site. Twelve retractable awnings (Gray *et al.*, 2016) were placed over the plots to block all water from any rainfall event in both wet and dry treatments. Drip irrigation was supplied once a week in order to maintain greater soil moisture in the wet treatment.

**Fig. 1.**
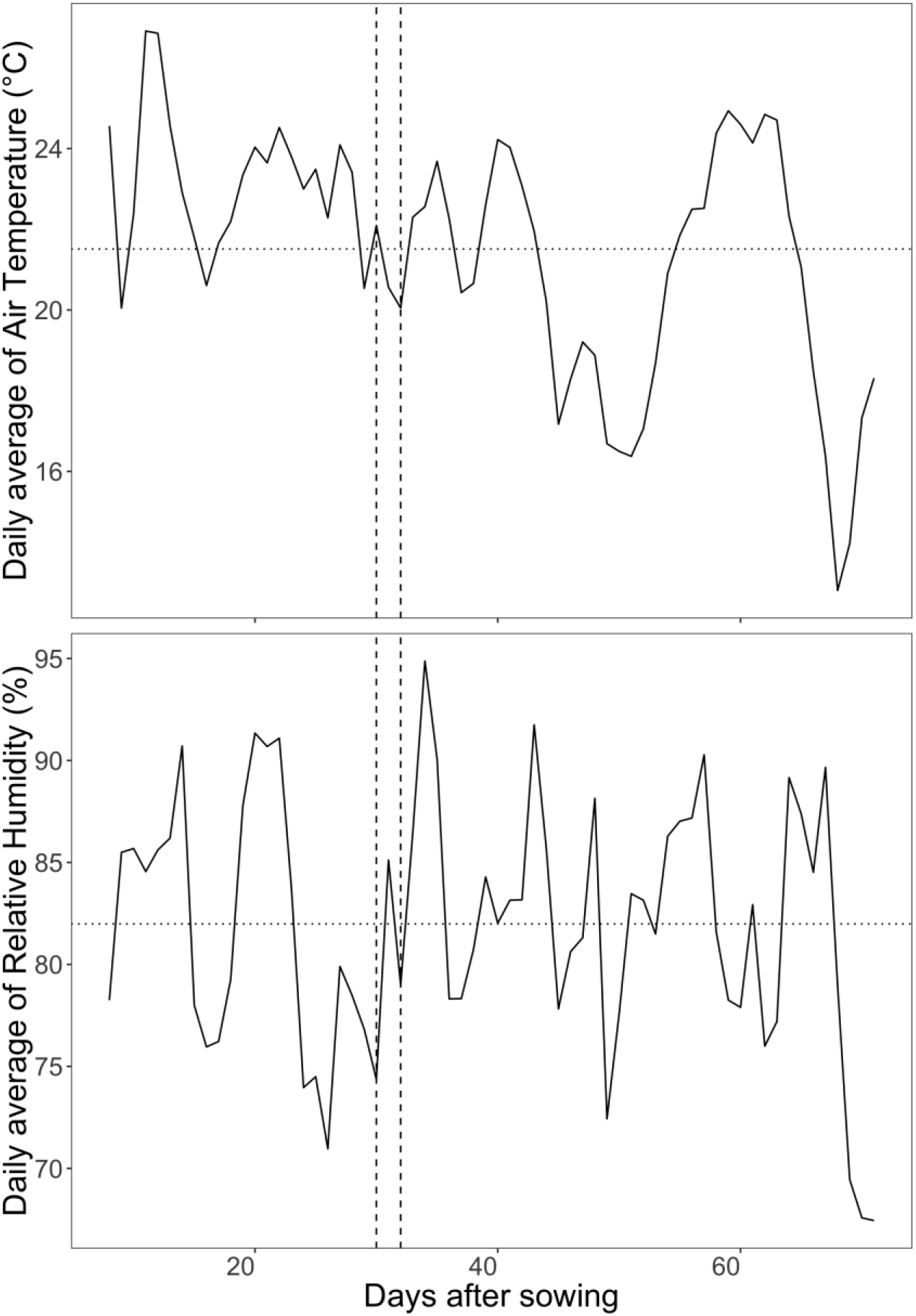
Daily average values of air temperature (a) and relative humidity (b) at SoyFACE experimental field site. The horizontal dotted line indicates the mean over the entire growing season. The vertical dashed lines indicate the days after sowing the canopy temperature measurements were collected in the field.

Each genotype subplot in the experiment measured 25 by 20 cm and contained 30 plants with a grid spacing of 5 cm between the plants. There was 25 cm space for the alleyway between two columns of plots and 10 cm spacing between the rows of plots. Each awning contained 66 subplots including six check plots of the B100 accession. The volumetric water content in the center of each awning was measured every 15 minutes throughout the growing season using soil moisture probes (CS650; Campbell Scientific) at 5 and 25 cm depths.

Canopy temperature of all field plots under both wet and dry treatments was measured 30 and 32 days after sowing (DAS) once canopy closure had occurred in all plots. A telescopic boom lift was used to collect images from a height of 9.1 m above the ground using a handheld infra-red camera (FLIR T400, FLIR Systems, Boston, MA, USA). On each date, one infrared and one RGB image was acquired for each awning, which consisted of 66 plots (Figure 2). The time of the measurements was between 11 am and 3 pm. Infrared imaging was performed only during clear and sunny weather conditions. Data from the 36 pixels at the center of each genotype subplot was used to estimate the canopy temperature (FLIR Tools, FLIR Systems, Boston, MA, USA). This ensured that temperature data were only sampled from pixels completely covered by plant canopy and not containing data from soil in the nearby alleys between plots.

**Fig. 2.**
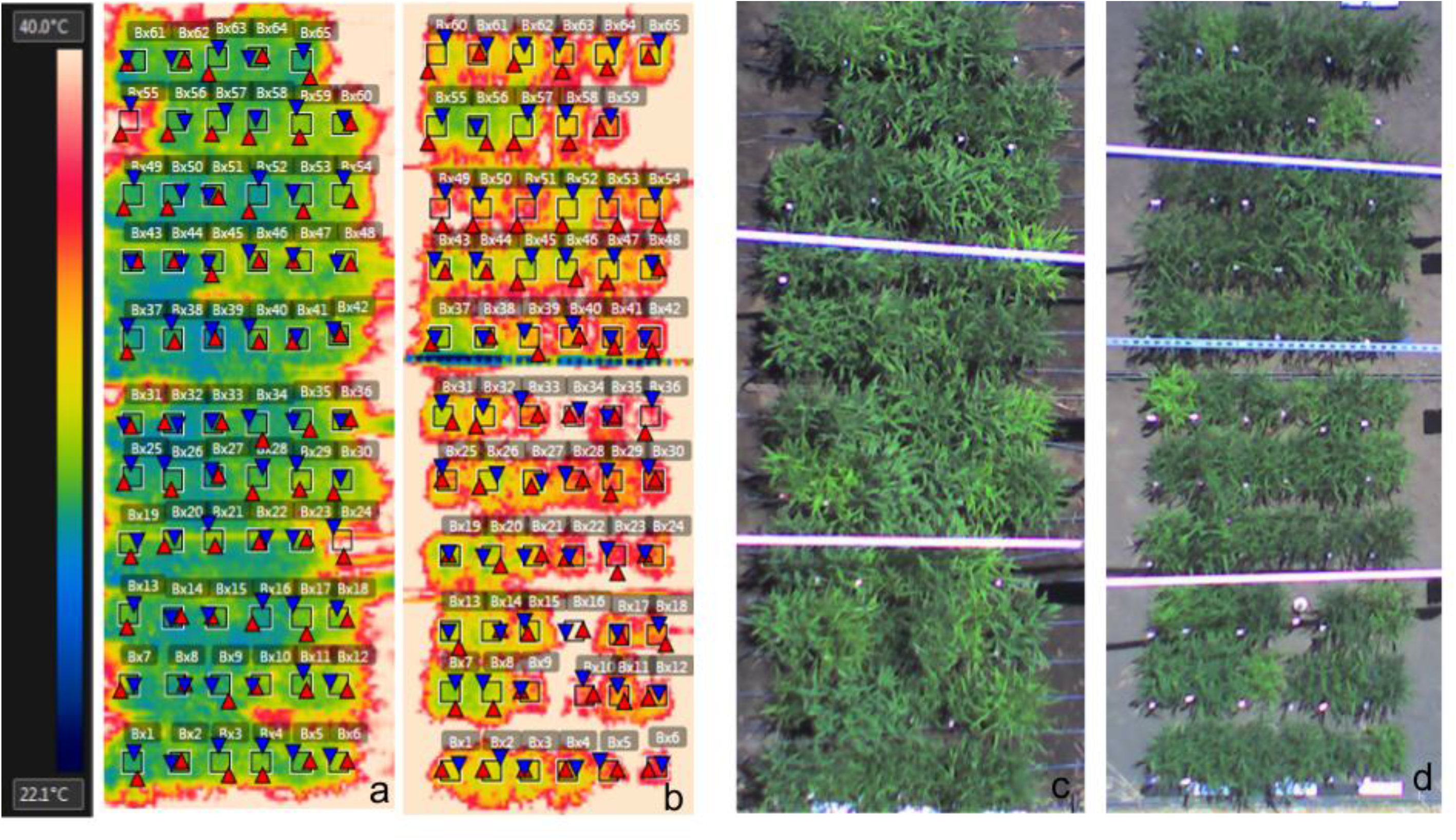
Aerial infra-red and RGB images of Setaria subplots under awnings in wet and dry treatments. Infra-red image of wet awning (a) and dry awning (b). RGB image of wet awning (c) and dry awning (d). The square boxes are the measured area of each subplot canopy.

Three plants from the center of each plot were destructively harvested 30 days after panicle emergence to estimate the shoot biomass. The plants were cut at the base and the leaf, stem and the panicles were separated and dried at 65°C. The dried weights of leaf, stem and panicle were summed to obtain the total shoot biomass.

### Data analysis

The greenhouse experiment was conducted with four replicates of each RIL arranged in a randomized complete block design with 120 genotypes as described in the equation below, where Y _ij_ is the individual observation of the trait of interest, μ is the overall mean, Genotype _i_ is the effect of the i^th^ genotype, Block _j_ is the effect of the j^th^ block and ε _ij_ is the error term.

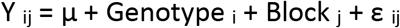

The field experiment was conducted as a randomized complete block design in a split plot arrangement with 3 blocks, 2 treatment conditions, 12 awnings nested within treatments and blocks and 120 genotypes as described below

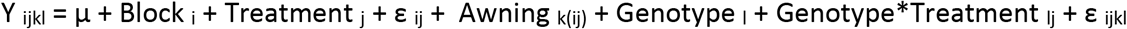

where Y _ijkl_ is the individual observation of the trait of interest, μ is the overall mean, Block _i_ is the effect of the i^th^ block, Treatment _j_ is the effect of the j^th^ treatment and ε _ij_ is the first error term, Awning _k(ij)_ is the k^th^ awning nested within Block _i_ and Treatment _j_, Genotype _l_ is the l^th^ genotype, Genotype*Treatment _lj_ is the interaction between Genotype l and Treatment j and ε _ijkl_ is the second error term.

The broad sense heritability on a line mean basis was computed using the variance components from the mixed model using the below formula.

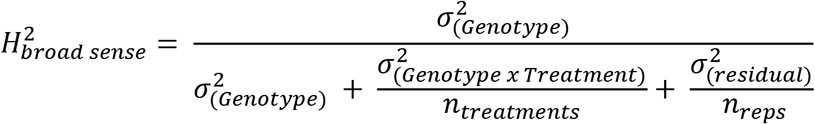

The variance components from the mixed model were extracted using lme4 package in R (Bates *et al.*, 2015). Best linear unbiased predictors (BLUPs) were calculated for each trait of interest using the experimental designs discussed earlier where genotypes and blocks were considered as random effects and treatment and awning as fixed effects.

The quantitative trail loci (QTL) mapping was performed on the BLUP values for stomatal density and canopy temperature under different treatments and sampling dates using ~1400 Single Nucleotide Polymorphism (SNP) markers. Mapping was performed using a custom biparental linkage mapping program (Feldman *et al.*, 2017) based upon the functionality encoded within the R/qtl (Broman *et al.*, 2003) and funqtl (Kwak *et al.*, 2014) packages in R. A two-step procedure was performed (Feldman *et al.*, 2017). First a single QTL model genome scan was performed using Haley-Knott regression to identify QTLs with LOD score higher than the significant threshold obtained through 1000 permutations at alpha 0.05. Second a stepwise forward/backward selection procedure was performed to identify an additive, multiple QTL model based upon maximization of penalized LOD score. The two-step procedure was conducted on all the traits and timepoints. QTLs that lie within 20 cM window are considered to be the same QTL.

## Results

### Soil moisture profile

Soil moisture content was equivalent in the wet and dry treatments at the beginning of the experiment (Figure 3). As time progressed, plants in the wet treatment continued to have adequate water supply (30 – 40 % vol/vol) throughout the growing period. By contrast, plants in the dry treatment experienced progressively drier soil conditions as the water they transpired was not replaced by rainfall or irrigation. The soil moisture was reduced in the dry treatment compared to the wet treatment at 5 cm and 25 cm depth by 20 DAS, resulting in a statistically significant interaction between treatment and time (p < 0.001) as well as significant overall effects of drought treatment (p < 0.001), depth (p < 0.001) and time (p < 0.001). Midday canopy temperature data was collected after this date, 30 and 32 DAS, when plants in the dry treatment were experiencing rapidly decreasing availability of soil moisture. This indicates that while plants in the dry treatment were subjected to limited water supply, they were still physiologically active i.e. drought stress was moderate.

**Fig. 3.**
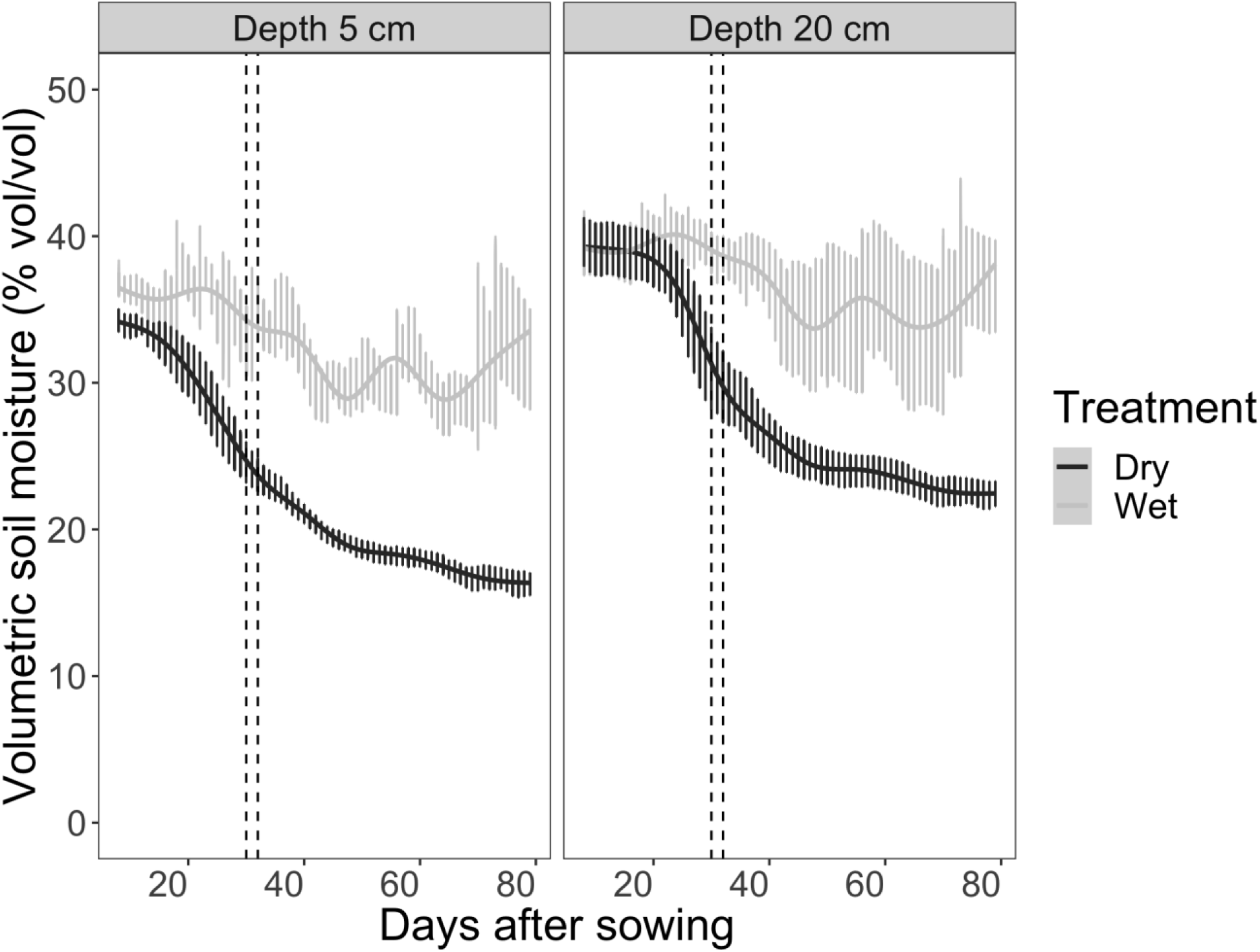
Soil volumetric water content (% vol/vol) at depths of 5 cm and 25 cm over the growing season in plots of Setaria supplied with either regular irrigation to maintain adequate water supply (wet treatment; light grey) or receiving no irrigation (dry treatment; dark grey). Rainfall was blocked from entering plots of both treatments using retractable rainout shelters. Data points and error bars shown the mean and standard error of three replicates per treatment. The dashed vertical lines indicate the dates when canopy temperature was measured.

### Genotypic variation in stomatal density and canopy temperature

Among the 120 RILs, stomatal density on the abaxial surface of the youngest fully expanded leaf ranged between 58 to 115 stomata/mm^2^ with a mean of 84 stomata/mm^2^ (Figure 4 and Figure 5). The broad sense heritability of stomatal density was 0.58. Among the 120 RILs, the mean canopy temperature at midday ranged from 28.8-31.9 °C at 30 DAS and 28.6-31.9 °C at 32 DAS in the wet treatment, and from 30.9-39.2 °C at 30 DAS and 29.3-38.1 °C at 32 DAS in the dry treatment. The mean midday canopy temperature across the RIL population was greater in the dry treatment than the wet treatment at both 30 DAS (32.9 °C versus 29.9 °C; p < 0.001) and 32 DAS (32.0 °C versus 29.6 °C; p < 0.001; Figure 6), with the treatment effect being slightly greater at 30 DAS (3.0 °C) than 32 DAS (2.4 °C). Midday canopy temperature was positively correlated between the two measurement dates for both wet (ρ = 0.78, p < 0.001) and dry (ρ = 0.66, p < 0.001) conditions, which gives confidence in the phenotyping method (Figure 7). The broad sense heritability of canopy temperature was 0.54 and 0.40 in 30 and 32 DAS, respectively.

**Fig. 4.**
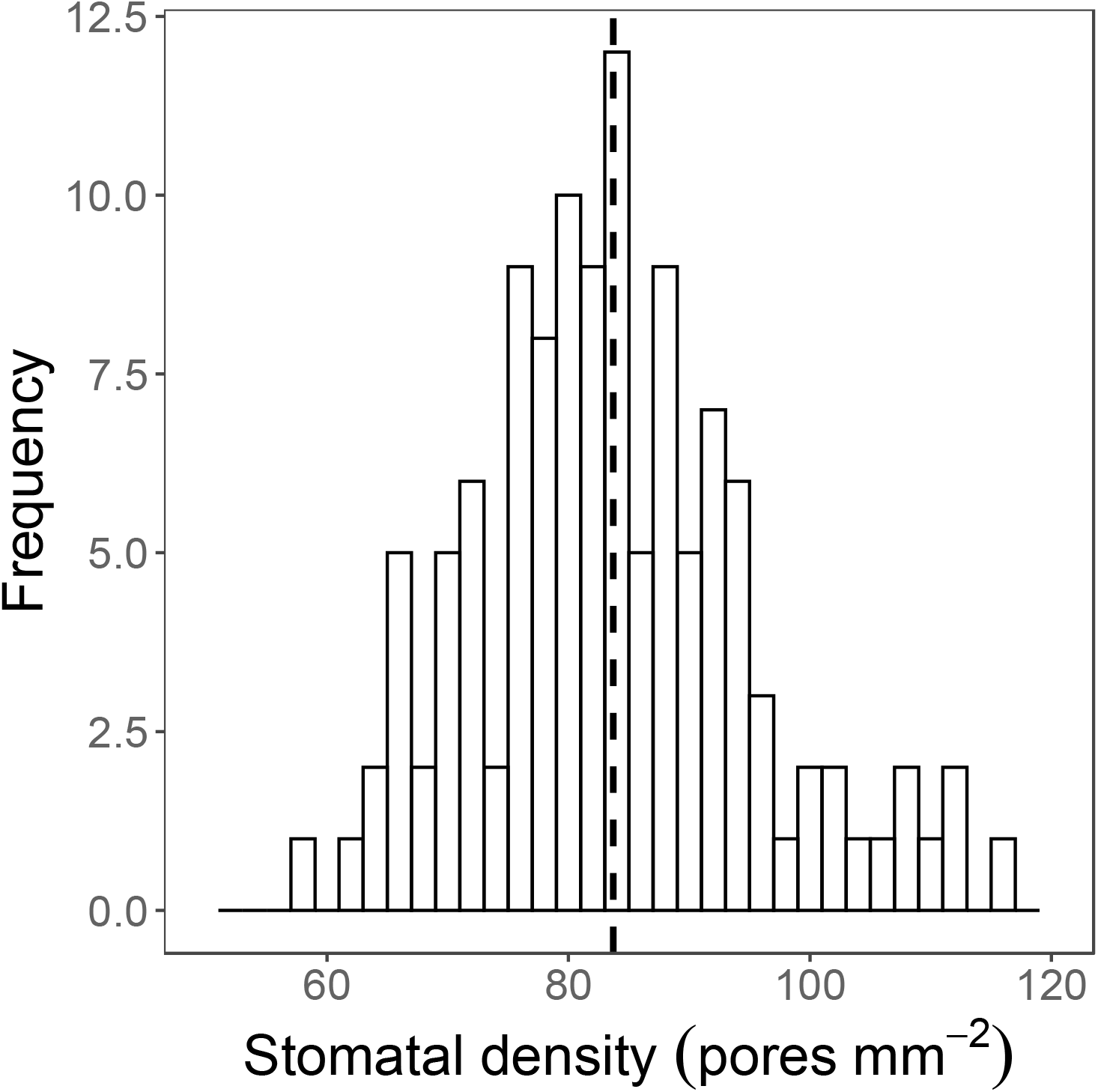
Frequency distribution of stomatal density (pores mm^−2^) of 120 recombinant inbred lines derived from a cross of *S. italica* and *S. viridis*, and B100 parental line. Data are genotype means derived from two fields of view per leaf from each of four replicate plants. The dotted vertical lines represent the population mean value.

**Fig. 5.**
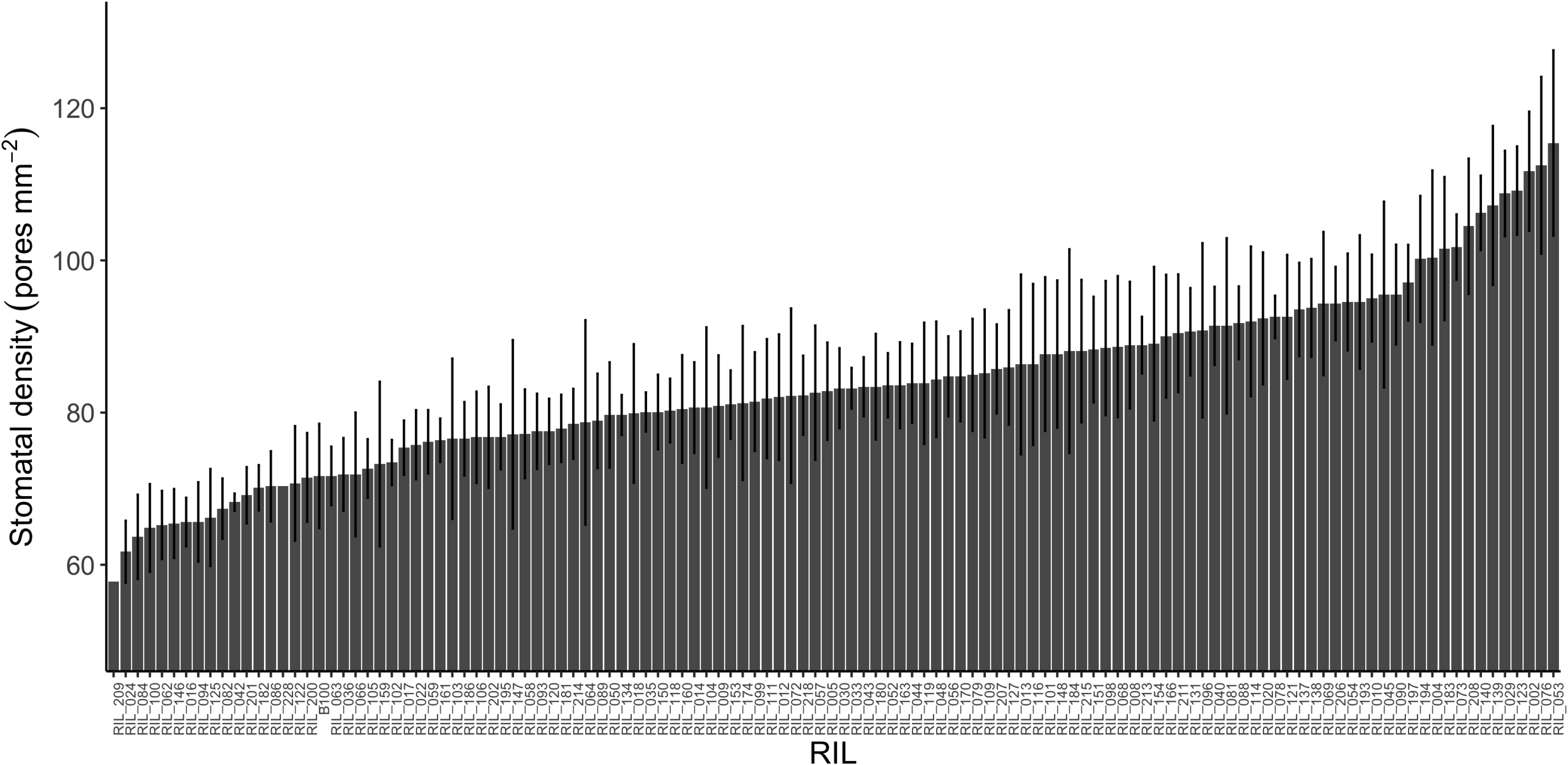
Stomatal density of 120 recombinant inbred lines derived from a cross of *S.italica* and *S.viridis, and B100 parental line.* Bars represent the genotype means (± standard error, n=4) derived from two fields of view from each of four replicate plants.

**Fig. 6.**
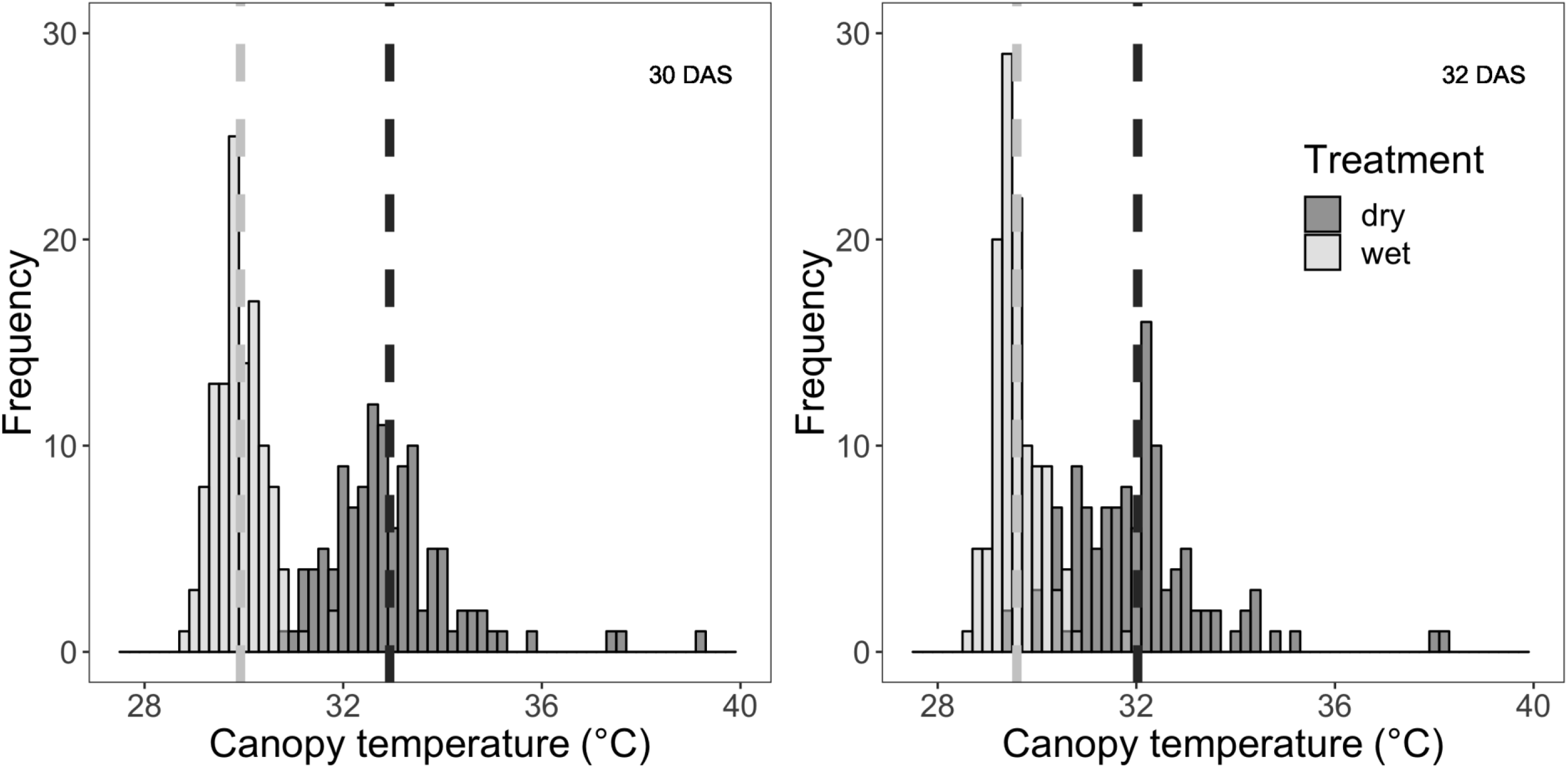
Frequency distribution of canopy temperature (°C) of 120 RILs in wet (light grey) and dry (dark grey) treatments at 30 and 32 days after sowing (DAS). Data are means derived from all pixels in the interior of three replicate plots per genotype. The dashed vertical lines represent the treatment mean value for each treatment.

**Fig. 7.**
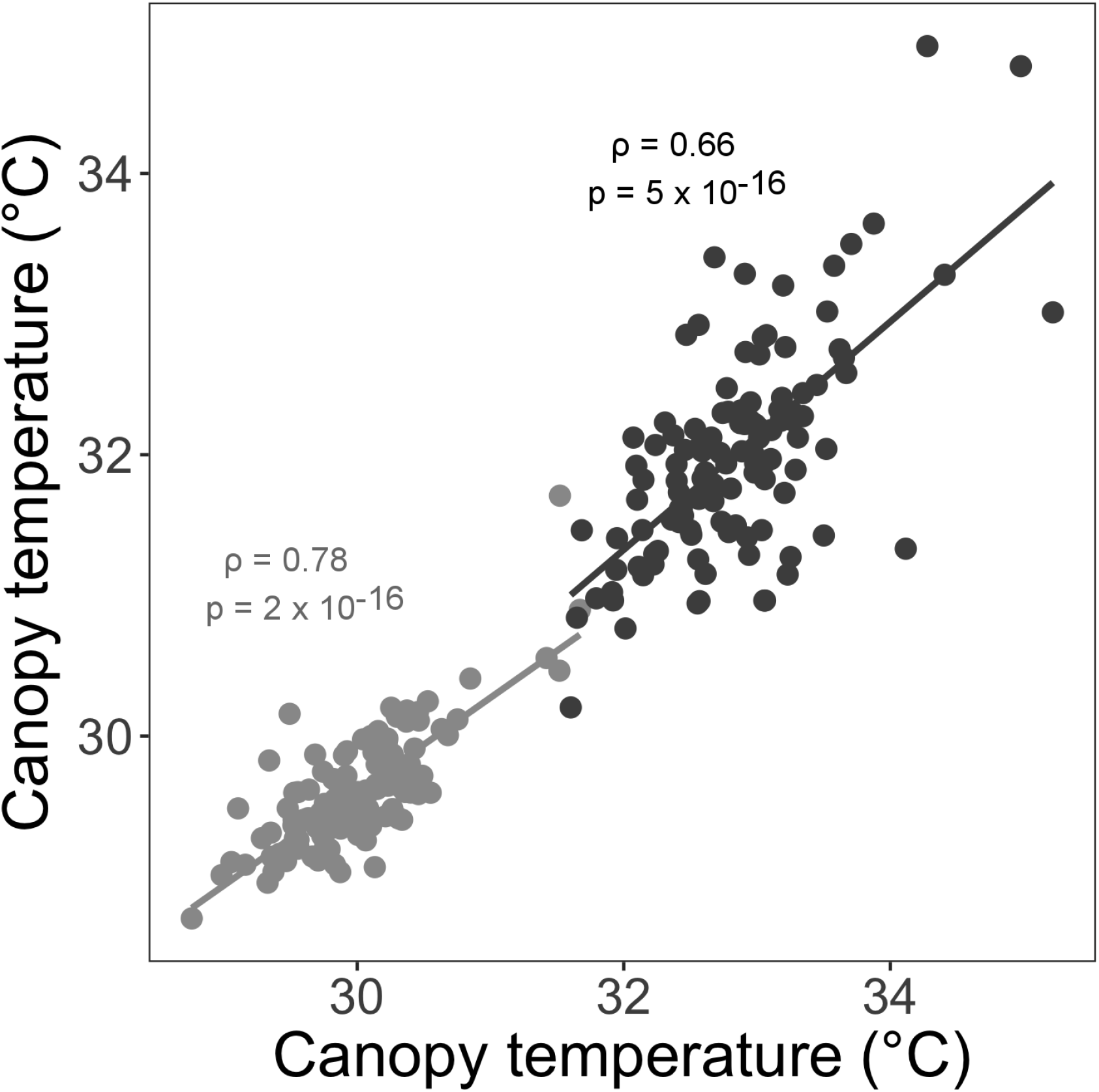
Scatterplot of midday canopy temperature for Setaria RILs and B100 on 30 DAS versus 32 DAS under wet 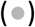 and dry treatments (●). Lines of best fit are shown along with the Pearson’s correlation coefficient (r) and associated p-value.

### Phenotypic relationships among canopy temperature, stomatal density and total biomass

Midday canopy temperature was negatively correlated with total above-ground biomass under both wet and dry treatments at both 30 DAS (wet: r = −0.38, p < 0.001; dry: r = −0.32, p < 0.001) and 32 DAS (wet: r = −0.49, p < 0.001; dry: r = −0.46, p < 0.001; Figure 8). The average increase in total above-ground biomass production associated with a decrease in midday canopy temperature of 1 °C was greater in the wet treatment than the dry treatment on both measurement dates (Table 1). The amount of variation in total above-ground biomass production explained by variation in midday canopy temperature was slightly greater in the wet treatment than the dry treatment on both sampling dates (Table 1). The parental line A10 recorded was one of the genotypes with lowest biomass and highest canopy temperature under both treatments and days of measurement, while the parental line B100 had trait values that were close to the mean of the population.

**Fig. 8.**
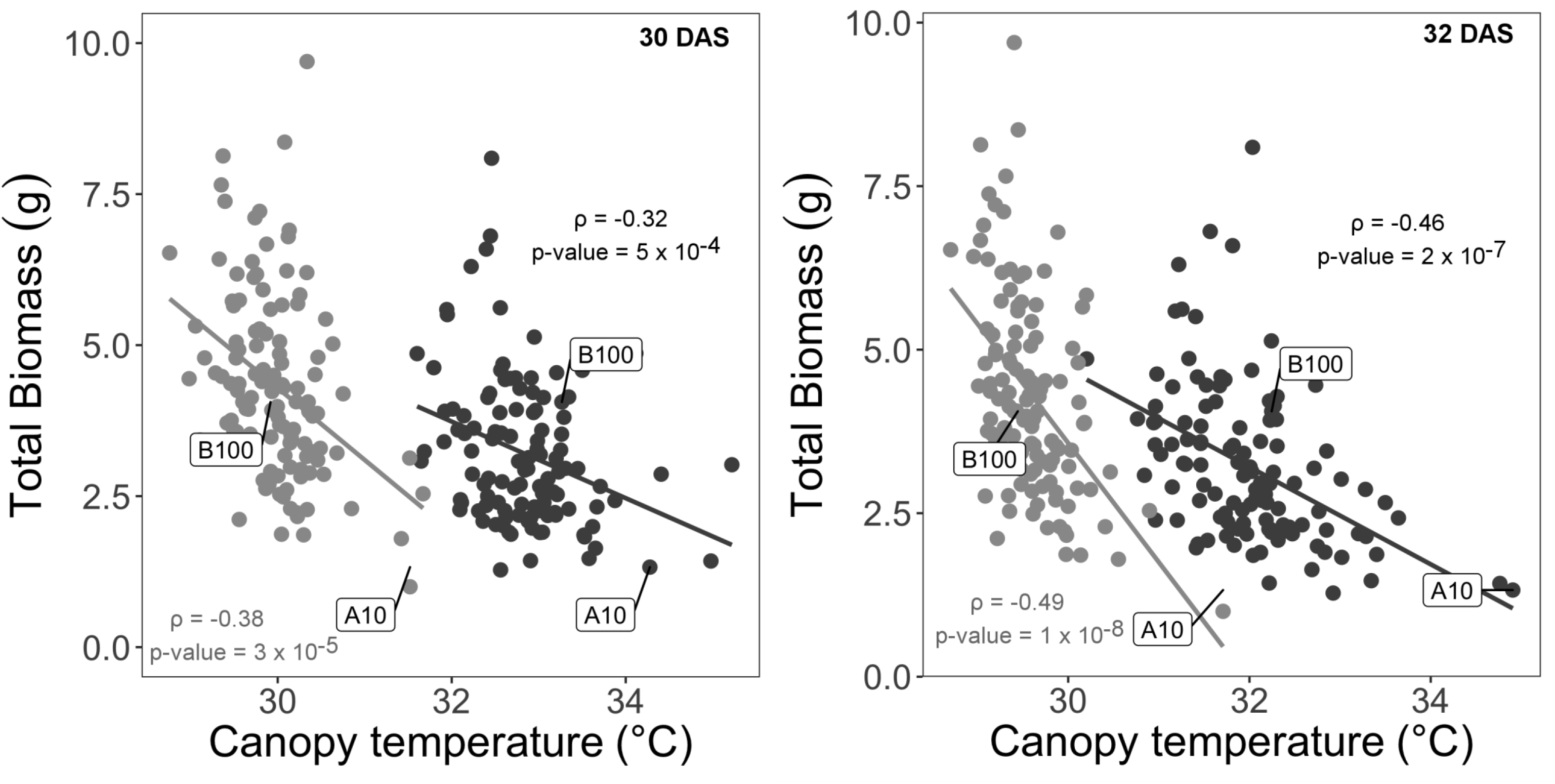
Scatterplot of total biomass (g per plant) in relation to canopy temperature (°C) for Setaria RILs and the parent lines (A10 and B100) under wet 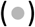 and dry conditions (●) at 30 and 32 days after sowing (DAS). Data are best linear unbiased predicted (BLUP) values for each genotype. Lines of best fit are shown along with the Pearson’s correlation coefficient (r) and associated p-value.

**Table 1.**
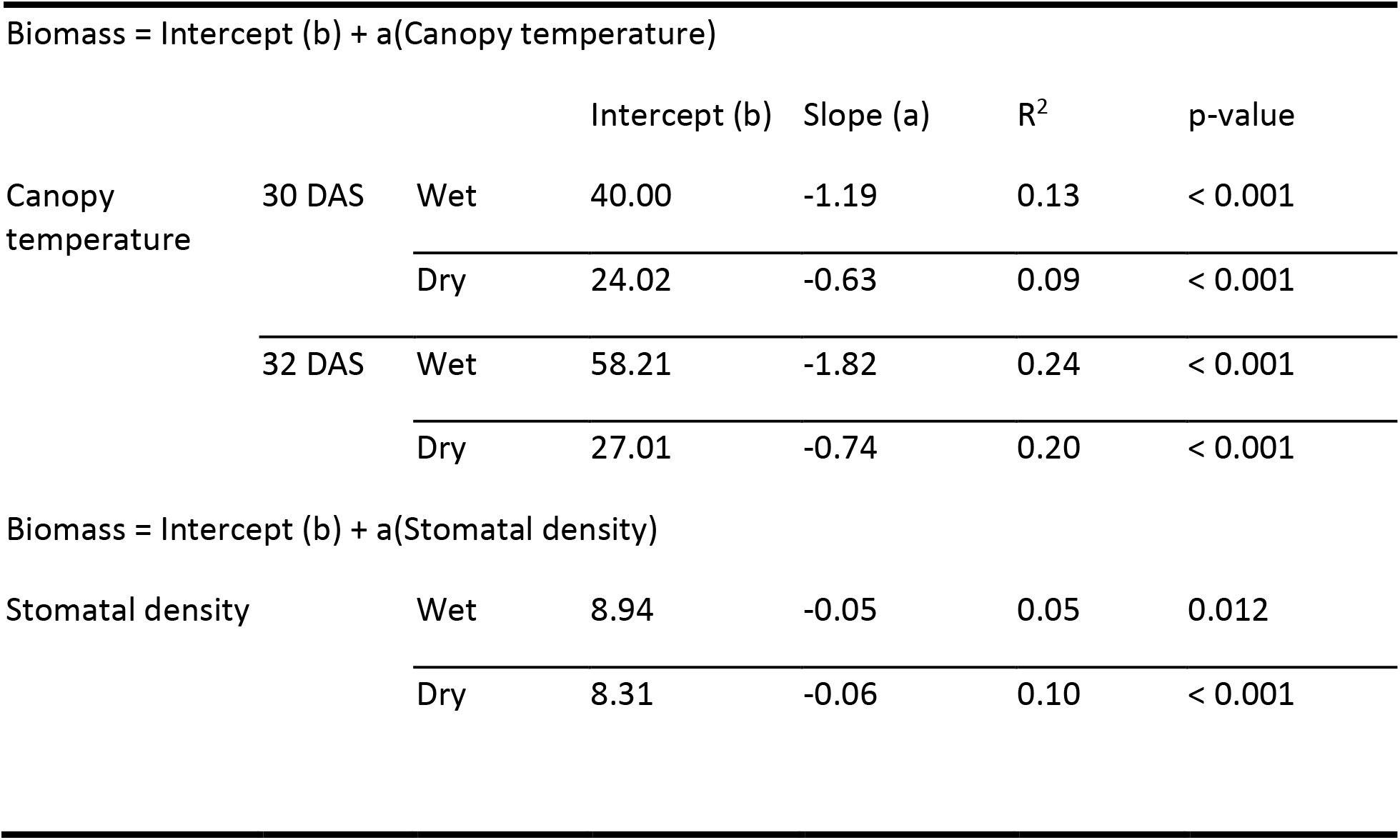
Regression parameters for total above-ground biomass (g per plant) in relation to canopy temperature (°C) and stomatal density (pores per mm^2^) of Setaria genotypes grown under wet and dry treatments.

Stomatal density was positively correlated with midday canopy temperature under both wet and dry treatments at both 30 DAS (wet: r = 0.40, p < 0.001; dry: r = 0.38, p < 0.001) and 32 DAS (wet: r = 0.37, p < 0.001; dry: r = 0.39, p = < 0.001; Figure 9). And, correspondingly, stomatal density was negatively correlated with total above-ground biomass under both dry (ρ = −0.33, p = < 0.001) and wet (r = −0.23, p = 0.012) conditions (Figure 10). The correlation between stomatal density and total biomass was stronger under the dry treatment than the wet treatment.

**Fig. 9.**
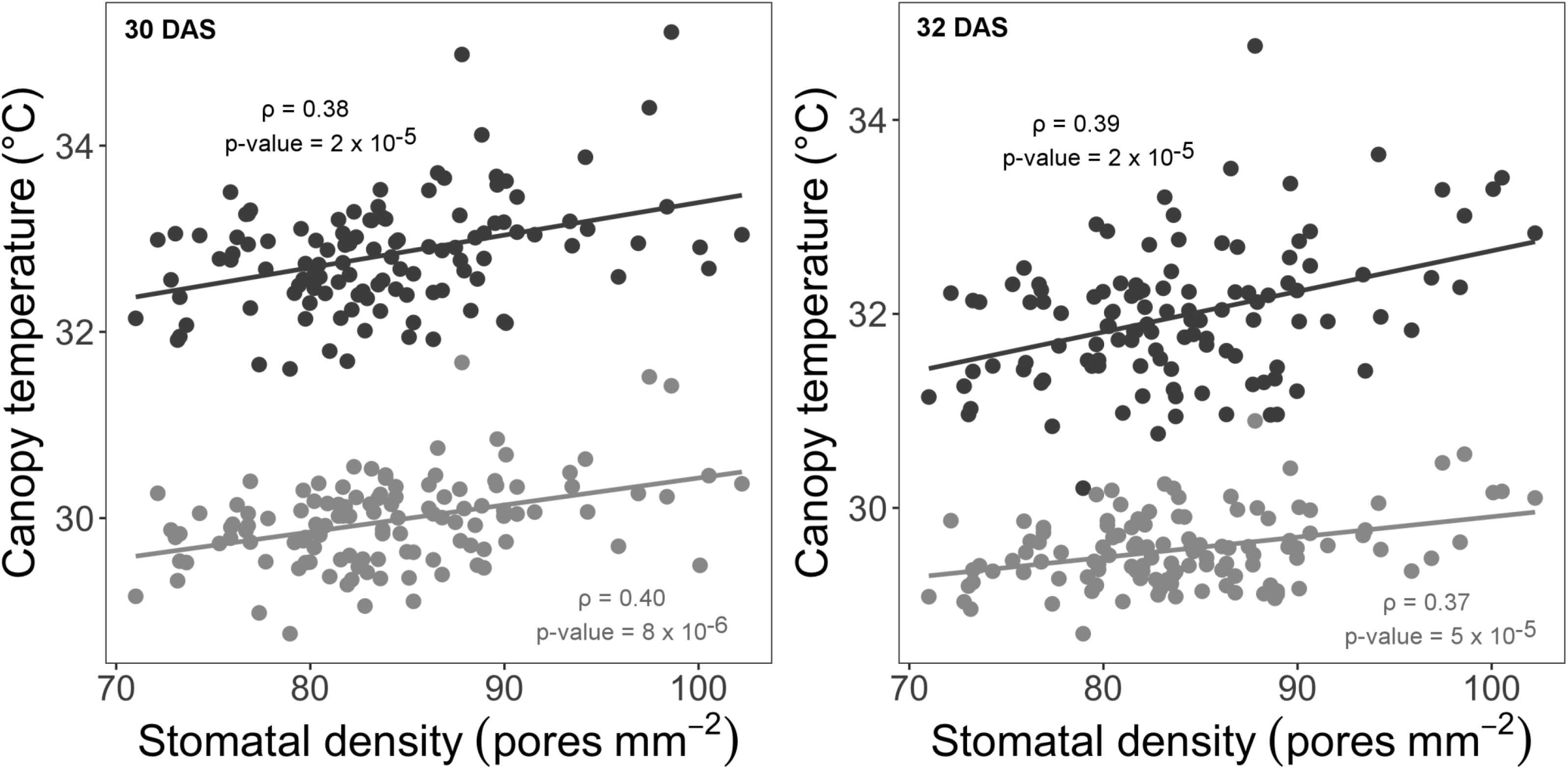
Scatterplot of canopy temperature (°C) in relation to stomatal density (pores mm^−2^) for Setaria RILs and the parent lines (A10 and B100) under wet 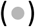 and dry (●) conditions at 30 and 32 days after sowing (DAS). Data are best linear unbiased predicted (BLUP) values for each genotype. Lines of best fit are shown along with the Pearson’s correlation coefficient (r) and associated p-value.

**Fig. 10.**
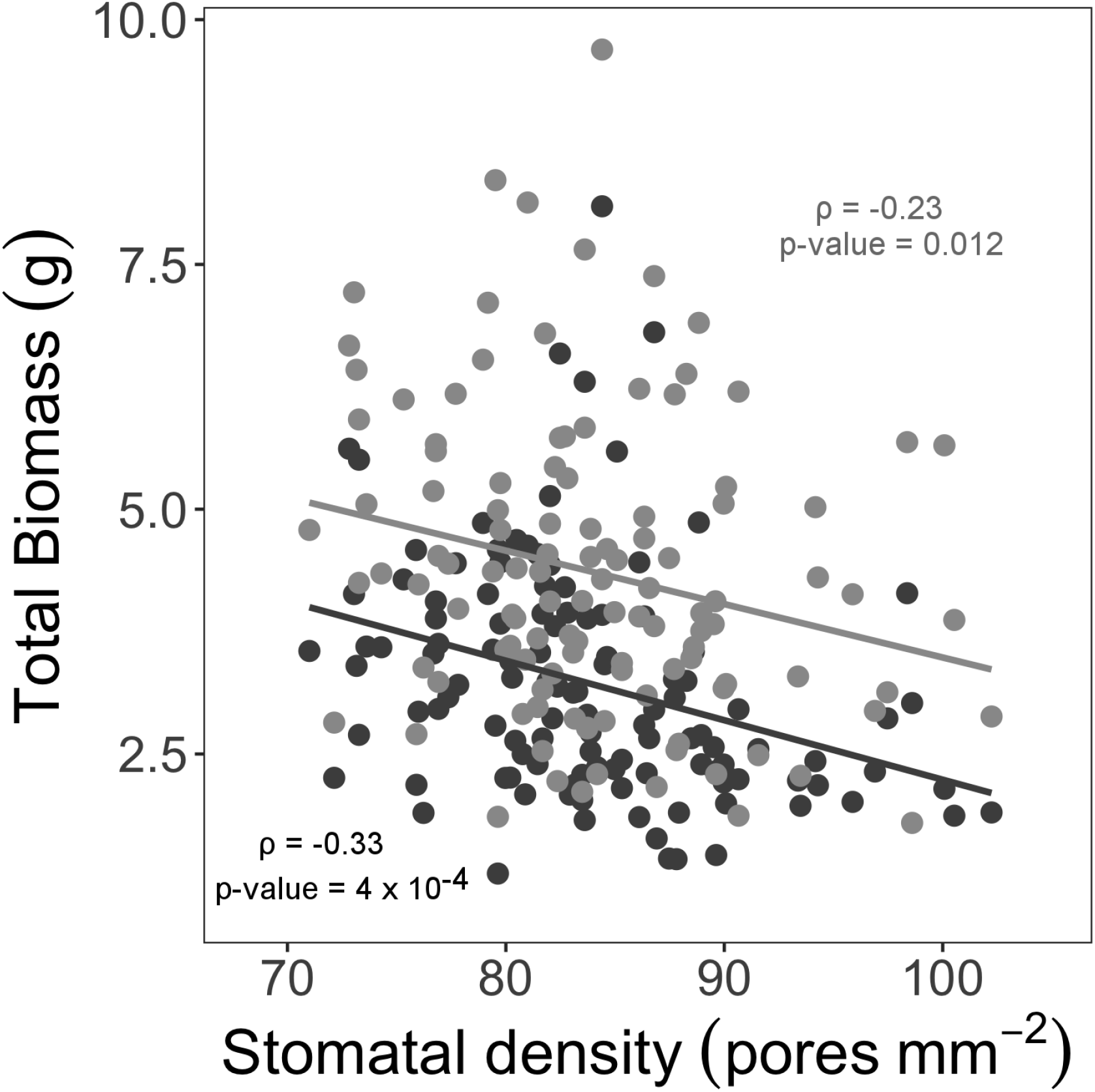
Scatterplot of total biomass (g per plant) relative to stomatal density (pores mm^−2^) for Setaria RILs and the parent lines (A10 and B100) under wet 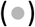 and dry (●) conditions. Data are best linear unbiased predicted (BLUP) values for each genotype. Lines of best fit are shown along with the Pearson’s correlation coefficient (r) and associated p-value.

### QTL mapping results

QTL analysis identified three significant loci for stomatal density and eight significant loci for canopy temperature (Table 2, Figure 11). The proportion of phenotypic variation associated with these QTLs ranged between 8 to 23 percent for both the traits. QTLs across different traits were considered to be overlapping if they were within a 20cM window and others that fall outside this window were considered to be unique QTLs (Feldman *et al.*, 2017). Two QTLs co-localized for both stomatal density and canopy temperature one on chromosome 5 and one on chromosome 9. All four alleles had negative additive effects, indicating that the B100 allele was reducing both stomatal density and canopy temperature.

**Table 2.** Putative quantitative trait loci (QTLs) for stomatal density and canopy temperature traits in the 120 F_7_ recombinant inbred line population derived from a cross of *S.italica* and *S.viridis,* and B100 parental line.

| Trait | Peak marker | Chr | Pos (cM)<sup>1</sup> | LOD at Peak<sup>2</sup> | Variance (%)<sup>3</sup> | Additive effect | Left CI (cM)<sup>4</sup> | Right CI (cM) |
| --- | --- | --- | --- | --- | --- | --- | --- | --- |
| SD | S5_42996052 | 5 | 104.8 | 8.3 | 20.8 | -3.8 | 101.1 | 106.6 |
|  | S9_10073675 | 9 | 45.6 | 5.0 | 11.6 | -2.3 | 40.4 | 52.7 |
|  | S9_50690449 | 9 | 136.5 | 3.7 | 8.3 | -2.0 | 133.0 | 146.9 |
| CT wet 30 DAS | S5_39309008 | 5 | 93.8 | 4.4 | 10.0 | -0.2 | 76.2 | 104.1 |
|  | S7_31494503 | 7 | 93.3 | 9.2 | 23.1 | 0.3 | 89.3 | 101.9 |
|  | S9_6724364 | 9 | 34.9 | 8.8 | 21.8 | -0.2 | 33.9 | 38.6 |
| CT wet 32 DAS | S7_31494503 | 7 | 93.3 | 3.8 | 12.0 | 0.2 | 89.3 | 101.9 |
|  | S9_6724364 | 9 | 34.9 | 6.4 | 21.0 | -0.2 | 32.8 | 38.6 |
| CT dry 30 DAS | S5_39309008 | 5 | 93.8 | 5.7 | 14.1 | -0.2 | 92.8 | 100.2 |
|  | S7_32133319 | 7 | 99.9 | 8.0 | 21.1 | 0.4 | 92.5 | 101.9 |
|  | S9_7218054 | 9 | 35.9 | 6.0 | 15.0 | -0.2 | 32.8 | 38.6 |

<sup>1</sup>Position of the peak marker in centimorgan (cM); <sup>2</sup> Logarithm of odds (LOD) of the peak marker, <sup>3</sup> Percentage of phenotypic variance explained by the QTL, <sup>4</sup> Left confidence interval of the QTL.

**Fig. 11.**
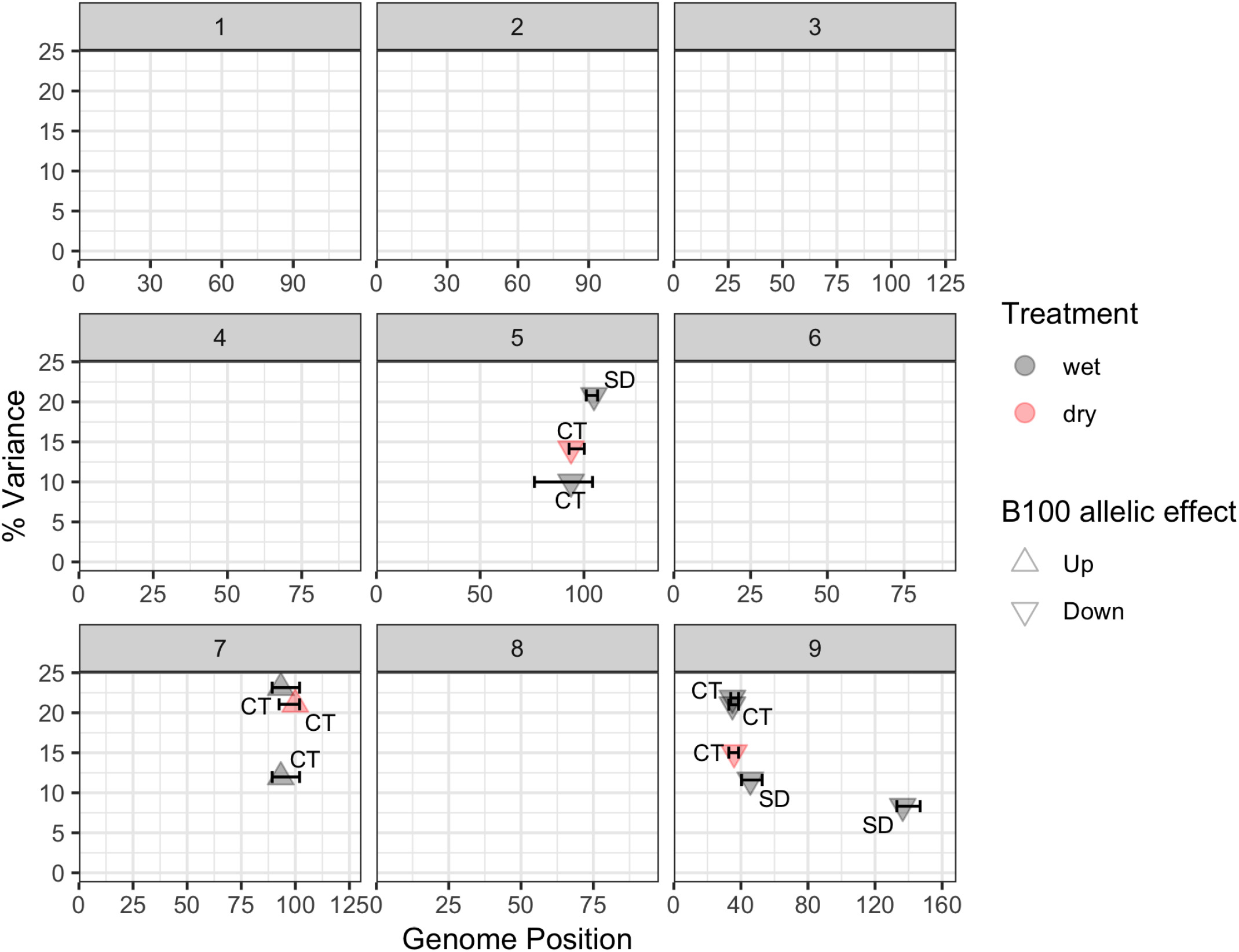
QTLs identified for stomatal density (SD) and canopy temperature (CT) under wet (grey) and dry (pink) treatments in the Setaria RIL population. Each panel corresponds to a chromosome. The arrow marks indicate the direction of the B100 allelic effect.

## Discussion

This study successfully characterized phenotypic and genetic variation in stomatal density and rates of canopy water use in Setaria, which can be used as a foundation for future studies to apply systems biology approaches to advance understanding of WUE and drought resistance in C_4_ species. Significant trait correlations were detected among stomatal density, canopy temperature and total above-ground biomass both in the wet and dry treatments.

The stomatal densities of RILs in this population (58 – 115 mm^−2^) were slightly greater than previously reported for faba bean (30 – 75 mm^−2^ Khazaei *et al.*, 2014) and wheat (36 – 92 mm^−2^ Schoppach *et al.*, 2016; 43 – 92 mm^−2^ Shahinnia *et al.*, 2016), but generally lower than Arabidopsis (90 – 210 mm^−2^ Dittberner *et al.*, 2018) and rice (273 – 697 mm^−2^ Laza *et al.*, 2010; 200 – 400 mm^−2^ Kulya *et al.*, 2018). While the magnitude of variation in stomatal density among the RIL population was sufficient to allow for QTL mapping and analysis of trait correlations, the parents of the population were not selected on the basis of this trait. Thus, the resulting magnitude of variation across the population was relatively modest. It would be valuable to investigate how much more variation for stomatal density may be found among genotypes within either *S. italica* or *S. viridis*, as well as the genus as a whole. The present study provided a proof of concept for the use of optical tomography to image the leaf epidermis. As proposed by Haus *et al.* (2015), optical tomography does not require sample preparation steps and can also be used on frozen leaf samples. This was significantly less laborious and more convenient than standard methods of taking leaf imprints of fresh leaves with dental gum and nail varnish (Rowland-Bamford *et al.*, 1990).

The magnitude of variation in canopy temperature across the Setaria RIL population was similar to that observed for sorghum (Awika *et al.*, 2017) and wheat (Mason *et al.*, 2013) RIL populations. Variation in canopy temperature among the RIL population were similar on 30 DAS (wet 3.1 °C, dry 8.3 °C) and 32 DAS (wet 3.3 °C, dry 8.8 °C) and canopy temperature was correlated across the two dates sampled for both the wet (r = 0.78) and dry treatments (r = 0.66). This might be considered surprising given the highly dynamic nature of canopy temperature in response to wind gusts, diurnal variation in solar radiation, and daily or seasonal variation in climate. But, the reproducibility of the data across dates is consistent with the comprehensive analysis by Deery *et al.* (2019), which analyzed 98 independent timepoints of canopy temperature data collected for a wheat population over 14 dates in two years. They concluded that canopy temperature could be reliably screened from one or two sampling points if data was collected under clear sky conditions in the afternoon, as was done in the current study. The present study also highlighted that Setaria as a highly tractable model for field trials because its small stature allows non-destructive, remote-sensing approaches to phenotyping, such as thermal imaging, to be performed on hundreds of replicated plots using hand-held cameras and a boom lift. This is significantly simpler in terms of data acquisition and data analysis than using drones or vehicles to gather data across field trials of crops with larger stature that require field trials covering larger areas (Deery *et al.*, 2016; Sagan *et al.*, 2019).

Canopy temperature was negatively correlated with the total above-ground biomass of the Setaria RILs under both wet and dry conditions. This is consistent with RILs that had higher temperatures due to less evaporative cooling being able to assimilate less CO_2_, and therefore producing less biomass, which was expected based on theory and previous studies (Fischer *et al.*, 1998; Jones, 2004). In addition, canopy temperature was significantly greater in the dry treatment compared to the wet treatment, which was consistent with stomatal closure reducing water use and evaporative cooling when there is limited water availability (Turner *et al.*, 2001). The relationship between canopy temperature and biomass was stronger in the wet treatment than the dry treatment on both measurement dates. This was reflected in canopy temperature explaining a greater proportion of variation in biomass (i.e. greater correlation coefficient) and a greater loss of biomass production per unit increase in canopy temperature under wet than dry conditions. This pattern of response is also consistent with prior observations (Bennett *et al.*, 2012; Mason *et al.*, 2013), but does not appear to have been the subject of much discussion. While it may seem initially counterintuitive that the relationship between the rate of water use and productivity would be weaker when water is limiting, it is consistent with genotypes that have inherently high rates of transpiration (i.e. cooler canopies) having greater reductions in productivity in response to drought stress than genotypes with inherently low rates of transpiration (i.e. warmer canopies). We suggest that this differential response may be conserved. And, it adds weight to the argument that genetic variation in WUE is best screened under well-watered conditions (Leakey *et al.*, 2019).

The positive correlation of stomatal density with the canopy temperature under drought stress suggests that the relationship between these two traits is complicated, since – if all else is equal – greater stomatal density would be expected to increase transpiration and lead to canopy cooling. Consistent with that theory, previous studies have reported that stomatal density is positively correlated with WUE (Xu and Zhou, 2008). However stomatal conductance is influenced by multiple factors, including stomatal density, maximum size and operating aperture (Dow and Bergmann, 2014; Faralli *et al.*, 2019). This implies that greater stomatal density within this population of Setaria RILs was associated with a developmental or functional shift that led to smaller stomatal apertures and lower rates of transpiration. As a result, within this population, lower stomatal density was also associated with greater biomass production. But, it should be noted that this relationship may be a function of the forced recombination across many parental alleles that is found in a RIL population. Breaking up gene linkage that can result from selection has been proposed to be a powerful approach to understand the biophysical basis for phenotypic relationships (Des Marais *et al.*, 2013). The observed positive correlation may reflect the developmental trade-off where stomatal size and stomatal density are widely found to be negatively correlated due to a limited amount of space on the epidermis (Shahinnia *et al.*, 2016; Faralli *et al.*, 2019), but this needs to be confirmed experimentally. By contrast, stomatal density was either not correlated or weakly, positively correlated with yield in wheat grown under both well-watered and drought treatments (Khazaie *et al.*, 2011; Schoppach *et al.*, 2016; Shahinnia *et al.*, 2016; Faralli *et al.*, 2019). So, the balance of trade-offs between stomatal density and aperture may be different among different biparental mapping populations, if not more generally in Setaria versus wheat. It would be valuable to compare if the same phenotypic relationship is observed across other biparental populations within these species as well as across natural accessions of these crops.

This study identified three unique QTL each for stomatal density and canopy temperature. All three of the canopy temperature QTL were robust in terms of being observed in both the wet and dry treatments. In addition, the canopy temperature QTLs on chromosomes 5 and 9 co-localized with QTLs for stomatal density (Figure 11). Genetic fine mapping would be required to discount the possibility that there are two loci in linkage at those locations. But, the observed pattern could be the result of pleiotropy, where a single locus regulates both traits. And, this would be consistent with the consistent direction of the allelic effects as well as positive correlation between the two traits, as well as the theoretical expectation that stomatal patterning on the epidermis influences transpiration rates. In that case, the ability to detect the same QTL in a greenhouse screen of stomatal density as for canopy temperature in the field suggests that rapid controlled environment screening might be a tractable way to accelerate progress in understanding and manipulating epidermal patterning and WUE in Setaria. The small stature of Setaria makes it particularly amenable for that approach. More broadly, the proportion of phenotypic variation explained by the stomatal density QTLs in Setaria were also similar to those of faba bean (Khazaei *et al.*, 2014), rice (Laza *et al.*, 2010), and wheat (Shahinnia *et al.*, 2016; Wang *et al.*, 2016).

Previous studies have identified many QTLs for different morphological and physiological traits using the same RIL population in Setaria in both controlled environment and field experiments (Mauro-Herrera and Doust, 2016; Feldman *et al.*, 2017; Banan *et al.*, 2018; Feldman *et al.*, 2018; Ellsworth *et al.*, 2020). These include measurements of traits with direct relevance to this study such as WUE of biomass production (i.e. biomass production relative to water use, as assessed by image analysis and metered irrigation on a high-throughput phenotyping platform linked to a controlled environment chamber). Meta-analysis of all the studies (Figure 12) reveals that QTL for stomatal density and canopy temperature overlap with QTLs for WUE, δ^13^C (Ellsworth *et al.*, 2020), plant height, panicle emergence, and various measures of above-ground productivity (Feldman *et al.*, 2017; Banan *et al.*, 2018) on chromosomes 5, 7 and 9. It is noteworthy that the percentage of the phenotypic variance explained by these QTLs for stomatal density and canopy temperature was typically equal to, or greater than, for the other traits assessed to date. One explanation for this would be that these loci directly regulate traits related to stomatal function and then indirectly influence the other traits via effects on crop water use. There is no reason to think the experimental design used here result in any greater statistical power to detect genotype to phenotype associations than the other studies. However, additional experimentation where all traits are measured simultaneously is needed to test this notion definitively.

**Fig. 12.**
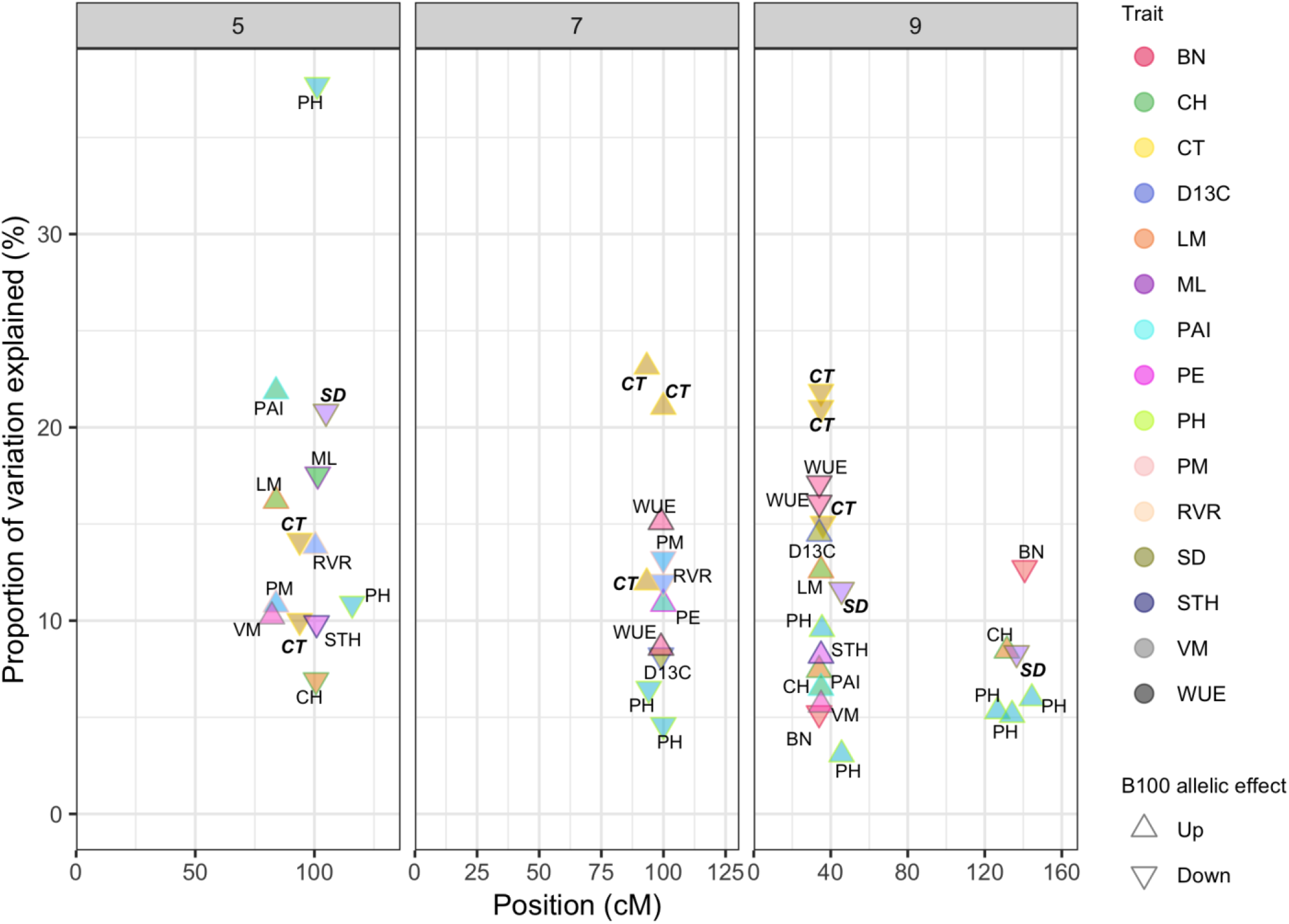
QTLs on chromosomes 5, 7 and 9 identified across multiple studies of *S. italica* ×*S. viridis* RIL population (Mauro-Herrera and Doust, 2016; Feldman et al., 2017; Banan et al., 2018; Feldman et al., 2018; Ellsworth et al., 2019). The arrow marks indicate the direction of the B100 allelic effect. The QTLs for stomatal density and canopy temperature identified in this study are denoted in bold and italics. BN – Branch number, CH – Culm height, CT – Canopy temperature, D13C – Delta13C, LM – Leaf mass, ML – Mesocotyl length, PAI – Plant area index, PE – Panicle emergence, PH – Plant height, PM – Panicle mass, RVR – Reproductive to vegetative mass ratio, SD – Stomatal density, STH – Secondary tiller height, VM – Vegetative mass, WUE – Water-use efficiency.

In conclusion, this study identified genetic loci in Setaria that are associated with variation in stomatal density as well as many other traits important to WUE, productivity and drought resistance. This suggests that Setaria is an experimentally tractable model system that would be highly suitable for more in-depth investigation of the mechanisms underpinning stomatal development and their influence on WUE in C_4_ species. An additional benefit to identifying QTLs and genes in Setaria is that it is also an agronomic crop, so the findings could have direct relevance to crop improvement programs as well as potentially translating into benefits for close relatives including maize, sorghum and sugarcane.

## Supplementary data section

Fig. S1. Field experiment layout for canopy temperature and biomass measurements

## Acknowledgements

Funded by the U.S. Department of Energy under Prime Agreement Nos. DE-SC0008769 and DE-SC0018277. We thank Dr. Timothy Wertin for helping with the stomatal density sample collection and other undergrads and summer interns for their help with field management. We also thank many project partners from the Danforth Plant Science Center, Carnegie Institute, Washington State University, and University of Minnesota that helped with transplanting seedlings.

## Author contributions

A.D.B.L. and I.B. conceived the original research plans. A.D.B.L., P.T.P., D.B., and R.E.P. supervised the experiments. P.T.P., collected the thermal images and processed the images. D.X. collected the stomatal images. P.T.P., D.B., and R.E.P., and L.F. managed the experiment and collected biomass data. P.T.P., M.F., I.B. and A.D.B.L. analyzed and interpreted the data. P.T.P. and A.D.B.L. wrote the article; M.F., I.B., D.B., R.E.P. and L.F. reviewed and commented on the article.

